# The role of immune inflammatory response and inflammatory infiltrates in major depression disorder and atopic dermatitis: new insights from machine learning

**DOI:** 10.1101/2023.10.15.562446

**Authors:** Han Jiang, Bizhen Gong, Mengxi Yao, Bo Pan, Hongya Zhang

**Author notes:** **Corresponding Author**: 1. Hongya Zhang, Department of dermatology, The First Affiliated Hospital of Anhui University of Chinese Medicine, HeFei, 230031, Anhui Province, China, Address: 117 Meishan Road, Hefei, Anhui Province, China,.; 2. Bo Pan, The First Affiliated Hospital of Naval Medical University, Shanghai, 200433, China, Address: 168 Changhai Road, Yangpu District, Shanghai, China,. Han Jiang, Bizhen Gong, Mengxi Yao contributed equally.

## Abstract

**Background:** Major depression disorder (MDD) and atopic dermatitis (AD) are distinct disorders involving immune inflammatory responses. This study aimed to investigate the comorbid relationship between AD and MDD and to identify possible common mechanisms.

**Methods:** We obtained AD and MDD data from the Gene Expression Omnibus (GEO) database. Differential expression analysis and the Genecard database were employed to identify shared genes associated with inflammatory diseases. These shared genes were then subjected to gene ontology (GO) and Kyoto Encyclopedia of Genes and Genomes (KEGG) pathway enrichment analyses. Hub genes were selected based on the protein-protein interactions using CytoHubba, and key regulatory genes were identified through enrichment analysis. Subsequently, we conducted immune infiltration and correlation analyses of the shared genes in AD. Finally, we employed three machine learning models to predict the significance of shared genes.

**Results:** A total of 17 shared genes were identified in the AD_Inflammatory_MDD dataset (S100A9, PTGER2, PI3, SNCA, DAB2, PDGFA, FSTL1, CALD1, XK, UTS2, DHRS9, PARD3, NFIB,TMEM158, LIPH, RAB27B, and SH3BRL2). These genes were associated with biological processes such as the regulation of mesenchymal cell proliferation, mesenchymal cell proliferation, and glial cell differentiation. The neuroactive ligand-receptor interaction, IL-17 signaling, and Rap1 signaling pathways were significantly enriched in KEGG analysis. SNCA, S100A9, SH3BGRL2, RAB27B, TMEM158, DAB2, FSTL1, CALD1, and XK were identified as hub genes contributing to comorbid AD and MDD development. The three machine learning models consistently identified SNCA and PARD3 as important biomarkers.

**Conclusion:** SNCA, S100A9, SH3BGRL2, RAB27B, TMEM158, DAB2, FSTL1, CALD1, and XK were identified as significant genes contributing to the development of AD and MDD comorbidities. Immune infiltration analysis showed a notable increase in the infiltration of various subtypes of CD4+ T cells, suggesting a potential association between the development of skin inflammation and the immune response. Across different machine learning models, SNCA and PARD3 consistently emerged as important biomarkers.

## 1. Introduction

Major depressive disorder (MDD) constitutes a chronic psychiatric ailment, afflicting over 300 million individuals worldwide(Pavlova & Ruda-Kucerova, 2023). Approximately 5% of adults globally endure the throes of MDD, making it a significant public health concern(Mpinga et al., 2023). MDD is characterized by neurochemical alterations within the cerebral domain. The latent life-related stressors associated with MDD are inclined to induce apoptotic events in mature neurons, incite immune responses, and diminish neurotrophic factors, thereby engendering anxiety and despondency(Lee & Jung, 2024). Atopic dermatitis (AD) is a chronic, inflammatory cutaneous malady, often accompanied by severe pruritus, which may exacerbate due to despondency, stress, or anxiety, thereby adversely affecting the life quality of the afflicted individuals(Abdelhadi et al., 2023). This ailment frequently occurs in the pediatric population and is associated with comorbidities of neuropsychiatric disorders,such as depression and anxiety(Wan et al., 2023). A profound interplay exists between the immune and nervous system functions in cutaneous physiology. The debilitating and chronically recurring lesions on the skin surface substantially impact the life quality of the afflicted individuals, and they may potentially lead to the onset of mental disorders(Iannone et al., 2022). Research has demonstrated that AD is an independent risk factor contributing to the onset of MDD, indicating a correlation between the occurrence of AD and MDD(Vinh et al., 2023).

Considerable research has been devoted to elucidating the interplay between the immune system and conditions such as MDD and AD(Ezzedine, Soliman, Li, Camp, & Pandya, 2023; Park, Jang, An, Choo, & Kim, 2020; van der Leek et al., 2020). Researchers have uncovered compelling links between MDD and the immune system. Notably, MDD patients often exhibit elevated serum levels of cytokines, such as the inflammatory markers C-reactive protein (CRP), interleukin-6 (IL-6), and tumor necrosis factor-alpha (TNF-α). Furthermore, studies have revealed the potential of anti-inflammatory medications, such as non-steroidal anti-inflammatory drugs and immunomodulatory agents, to ameliorate depressive symptoms in select individuals(Chan, Poller, Swirski, & Russo, 2023). Additionally, neuroinflammation and the gut-brain axis studies highlighted the significance of neuroimmune interactions in depression pathogenesis(Boufidou & Nikolaou, 2016). Furthermore, studies have clearly delineated the pivotal role of the immune system in the development of AD. AD patients typically exhibit aberrant activation of T cells and cytokines, including interleukin-4 and interleukin-13. Moreover, studies on the skin microbiome have unveiled a close nexus between immunodysregulation and AD, further enhancing our comprehension of this association(Boufidou & Nikolaou, 2016). Despite the substantial progress, the intricacies of the connections between depression, AD, and the immune system remain to be elucidated. Consequently, it is imperative to delve deeper into these relationships, particularly by employing advanced machine learning methodologies, to garner more profound insights.

The rapid advancement of machine learning and artificial intelligence has provided novel tools and methodologies for medical research(Fang et al., 2023). Machine learning excels in handling vast datasets, recognizing latent patterns, and predicting trends. The application of this technology holds promise in providing fresh insights, deepening our comprehension of these maladies, and aiding in unraveling the intricate relationships between the immune system, depression, and AD. The technology also deepens our understanding of these intricate interactions, unlocking new prospects for precision medicine and more efficacious therapeutic strategies(Greenfield, Feizpour, & Evans, 2023; Scodari, Chacko, Matsumura, & Jacobson, 2023).

## 2. Materials and methods

### 2.1 Data retrieval

We acquired the datasets for the MDD and AD from the Gene Expression Omnibus (GEO) database (https://www.ncbi.nlm.nih.gov/geo/)(Barrett et al., 2013). We downloaded the GSE168694 dataset from the GPL22120-25936 platform, and it contained the peripheral blood samples from 4 individuals with AD and 4 healthy controls(L. Yang et al., 2022). The GSE182740 dataset from the GPL570-55599 platform comprised skin tissue samples obtained from 26 subjects, including 20 patients diagnosed with AD and 6 healthy subjects. For MDD, we utilized the gene expression dataset denoted as GSE58430(Wang et al., 2015) downloaded from the GPL14550-9757 platform, and it consisted of 12 peripheral blood samples (6 patients with MDD and 6 healthy controls). GeneCards (https://www.genecards.org/)(Stelzer et al., 2016), a comprehensive database of human genomics and transcriptomics, was used to search for disease targets associated with inflammatory responses using the keyword “inflammatory.”

### 2.2 Identification of differentially expressed genes (DEGs)

The raw expression matrix was normalized and processed using R (4.2.2) software. The DEGs were screened from the GSE168694, GSE182740 and GSE58430 datasets using the “limma” R package(Y. Chen, Chen, & Lei, 2022), and the batch effect of the GSE168694 and GSE182740 datasets was eliminated using the R package “sva.” The identified DEGs were evaluated to ensure that they met the requirements of adj. P < 0.05 and |log2 (Fold-change)| > 1. The R package “sva” was used to eliminate the batch effect of the two AD datasets.

### 2.3 Acquisition and enrichment of immune-inflammatory response genes shared by AD and MDD

We employed Wayne’s Chart Tool of the Xiantao Love (https://www.xiantaozi.com) to extract the pertinent inflammatory genes denoted as ‘AD_inflammatory’ from the DEGs identified in AD. Subsequently, we compared the DEGs identified in MDD with the ‘AD_inflammatory’ to identify the overlapping genes, herein referred to as ‘AD_Inflammatory_MDD-related DEGs.’ These shared genes were then subjected to further scrutiny through functional enrichment analysis. The Gene Ontology (GO) enrichment analysis was executed using the ‘enrichplot’ and ‘ggplot2’ software packages within the R environment(Gustavsson, Zhang, Reynolds, Garcia-Ruiz, & Ryten, 2022). Metascape (https://metascape.org) was used to analyze the potential Kyoto Encyclopedia of Genes and Genomes (KEGG) signaling pathways. Statistical significance was set at P < 0.05.

### 2.4 Analysis of protein-protein interactions (PPIs) and identification of hub genes

We used the STRING database (https://string-db.org/) to elucidate the PPIs within the trio, establishing a minimum required interaction score of 0.4 while obfuscating disconnected nodes within the network. Following the analysis of node relationships in the STRING network graph via Perl, we imported the dataset into Cytoscape v3.7.1 and generated the intersection genes employing the built-in ‘cytohubba’ plugin(Luna, Shah, Sander, & Shannon, 2023). Subsequently, we exported the computation results into R and employed the R package “UpSetR” to score and rank hub genes(Conway, Lex, & Gehlenborg, 2017), ultimately obtaining the core intersection genes. Finally, we visualized the core intersection genes using the R package “pheatmap.”

### 2.5 Functional enrichment of core genes

We conducted GO function enrichment analysis for the core genes using the R package “GOplot” with a predefined statistical significance threshold of P < 0.05. Subsequently, we identified the top eight significantly enriched GO terms utilizing the “RColorBrewer” and “circlize” packages to elucidate the associations between genes and enriched functions.

### 2.6 Immune infiltration and correlation analyses

After the normalization process for immune infiltration analysis, we first used the R package “CIBERSORT” to analyze the corrected data of GSE168694 and GSE182740 microarrays. “CIBERSORT” estimates the proportion of different immune cell types via a complex mathematical model using known immune cell characteristics and relationships among the sample data. These immune cell types can include various T-cell subtypes, B-cells, macrophages, and others. Statistical significance was set at P < 0.05, and the immune infiltration analysis results were visualized using the R package “vioplot.” Subsequently, R package “limma” was used to analyze the correlation between the three shared genes, AD_Inflammatory_MDD, and the immune cell types in the immune infiltration analysis results.

### 2.7 Machine learning screening

We used three different machine learning models, Random Forest (RF), Support Vector Machine (SVM) and Generalized Linear Model (GLM), to predict the AD_Inflammatory_MDD triple shared genes. The training was performed on the joint AD two-chip dataset to determine which genes were most critical for predicting sample type. Using the R packages “randomForest,” “caret,” “DALEX,” and “kernlab,” we plotted gene importance graphs and generated the most important genes in each model.

## 3. Result

### 3.1 Identification of DEGs

After normalizing the combined AD datasets (Fig. 1), 637 DEGs were obtained, comprising 281 up-regulated and 356 down-regulated genes. For the MDD dataset, 474 DEGs were obtained, containing 198 up-regulated and 276 down-regulated genes (Fig. 2). Volcano plots showed the expression profiles of DEGs within the GSE168694, GSE182740, and GSE584303 datasets, as displayed in Fig. 4. Furthermore, heatmaps provided insights into the top 50 DEGs associated with both AD (Fig. 4a) and MDD (Fig. 4b).

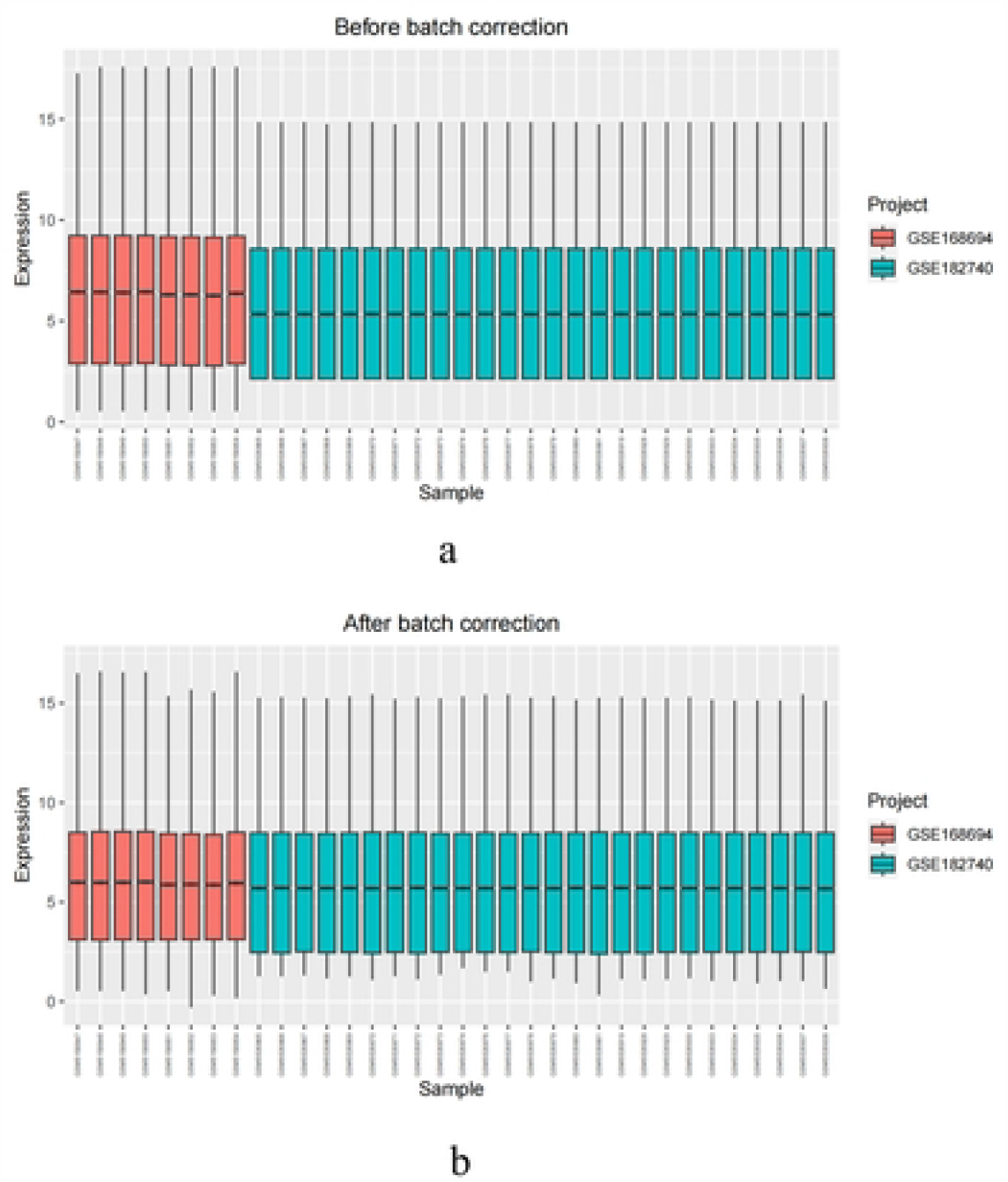

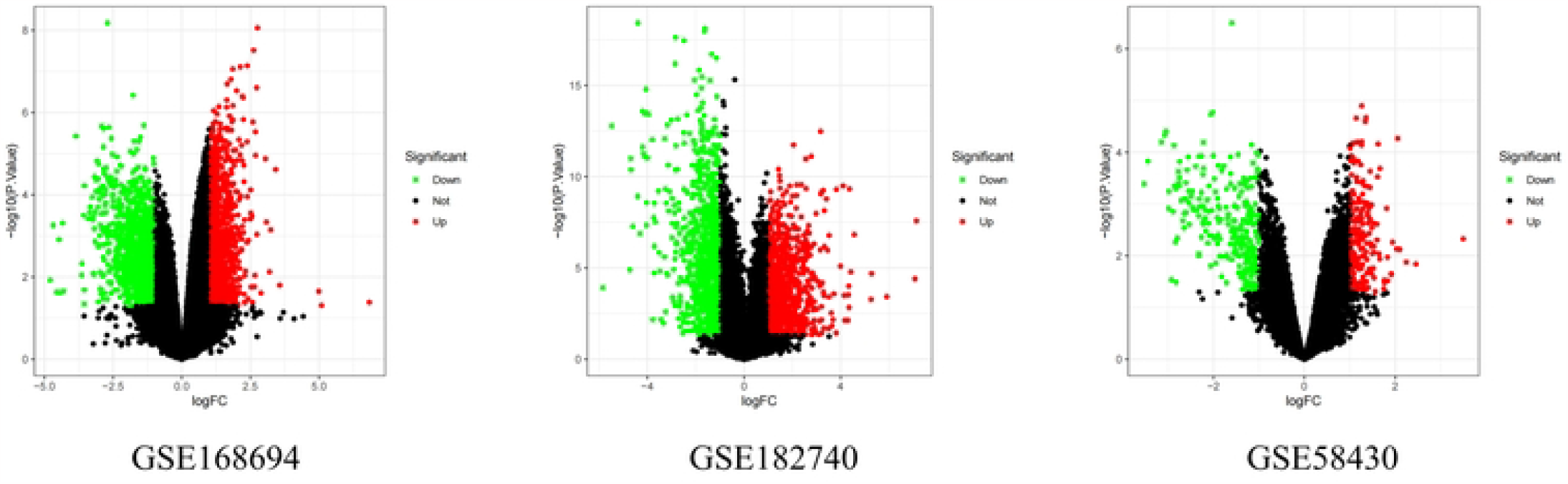

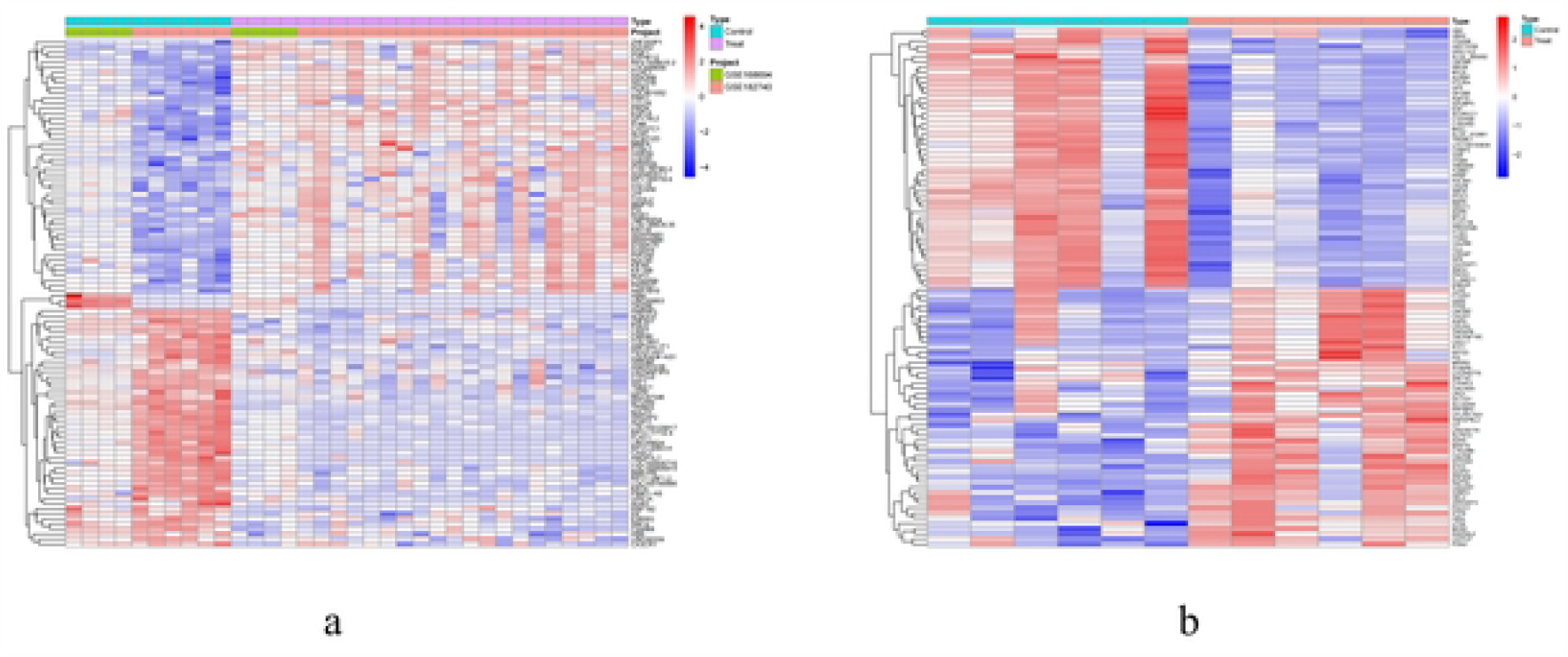

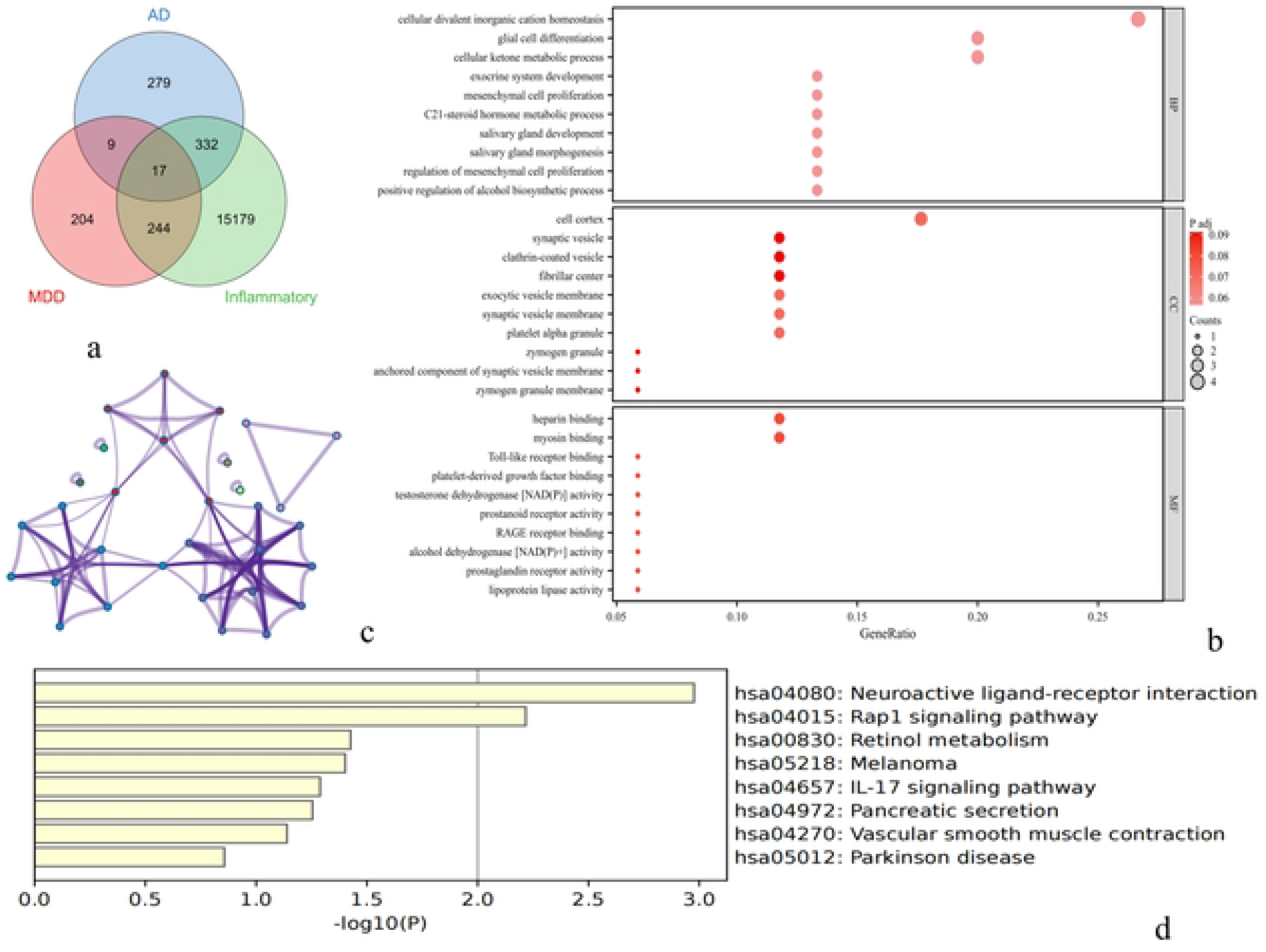

### 3.2 Identification of AD_Inflammatory_MDD functional enrichment genes

We obtained 17 shared genes (S100A9, PTGER2, PI3, SNCA, DAB2, PDGFA, FSTL1, CALD1, XK, UTS2, DHRS9, PARD3, NFIB, TMEM158, LIPH, RAB27B, and SH3BRL2) with the overlapping crossover between AD, MDD and inflammatory genes (Fig.4a). The GO and KEGG analyses of these 17 genes were performed to determine the regulatory and signaling pathways in which they are involved. As shown in Fig.4b, the genes were involved in biological processes that regulate mesenchymal cell proliferation, mesenchymal cell proliferation, cellular ketone metabolic processes, and glial cell differentiation. Cellular components involved in regulation were the cell cortex, platelet alpha granules and synaptic vesicle membranes. The molecular functions involved in regulation were Toll-like receptor binding, RAGE receptor binding and myosin binding. Moreover, the KEGG enrichment analysis showed (Fig.4c and 4d) that the signaling pathways that these genes were involved in were neuroactive ligand-receptor interaction, IL-17 signaling and Rap1 signaling pathways.

### 3.3 Identification and enrichment analysis of core genes

The STRING PPI mutual analysis of shared genes showed that there were 17 edges for 17 nodes (Fig.5a). The analysis of each node of the 17 PPI network showed (Fig.5b) that there are 2 up-regulated genes (PI3 and S100A9) 10 down-regulated genes (CALD1, FSTL1, DAB2, NFIB, PARD3, TMEM158, SNCA, RAB27B, SH3BGRL2 and XK). Nine hub genes were finally screened out (Fig.5c, Table 1), and their differential expression in the various AD datasets is shown in Fig.5d. Subsequent GO enrichment against the hub genes showed (Fig.5e) that they are mainly involved in biological processes such as positive regulation of secretion by cell, positive regulation of secretion, positive regulation of proteolysis, and synaptic vesicle recycling. The Circos analysis (Fig.5f) further demonstrated the relationship between function annotation and hub genes. The results showed that DAB2, RAB27B, SNCA, and S100A9 were closely related to the mentioned biological processes.

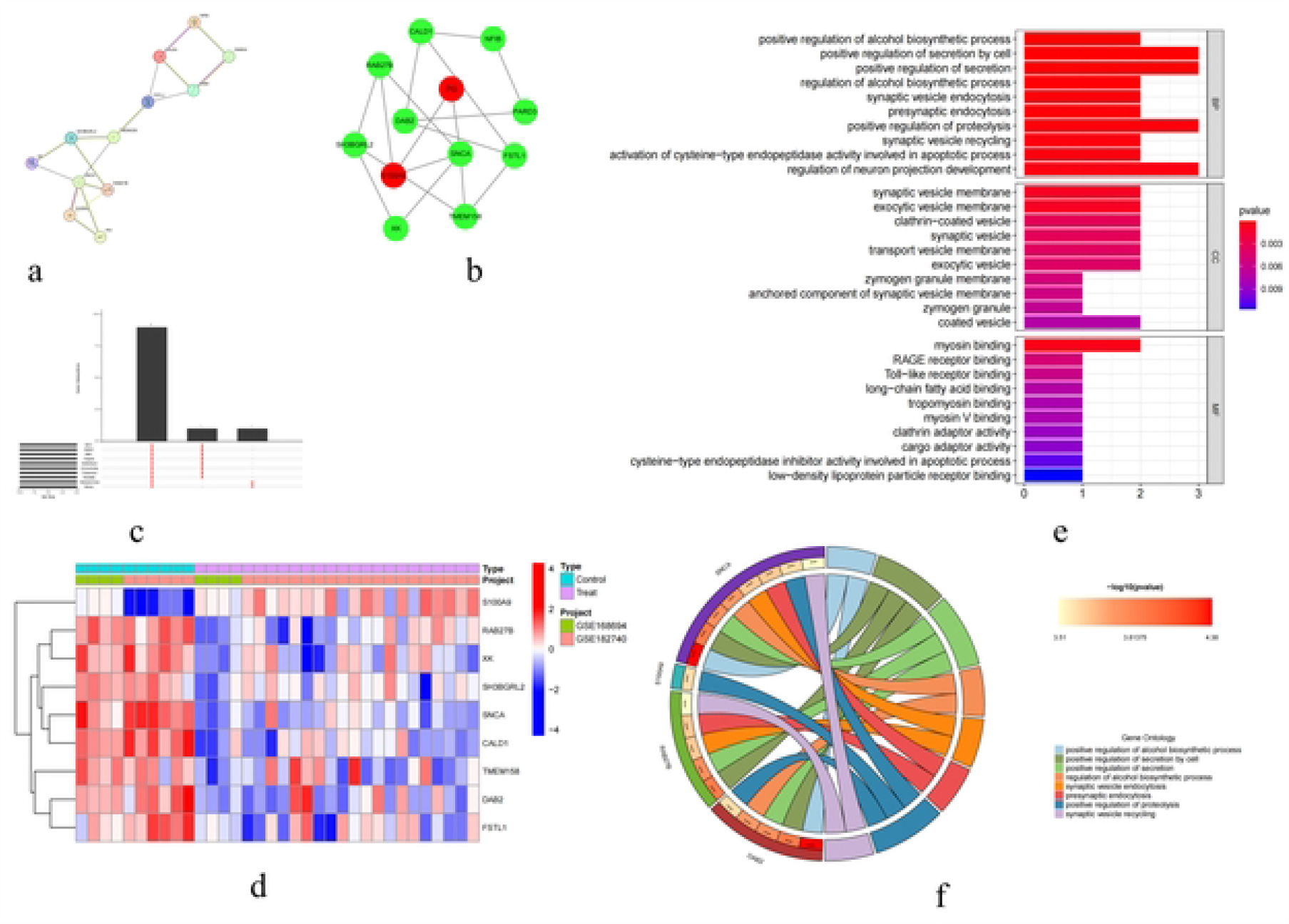

### 3.4 Results of immune infiltration analysis

We further investigated the immune infiltration differences in different samples and identified 22 types of immune cells in two joint datasets related to AD (GSE168694 and GSE182740). Fig.6a shows the distribution of the different types of immune cells. Furthermore, we found that the increased infiltration of resting memory CD4 T cells, activated memory CD4 T cells, follicular helper T cells, activated NK cells, and Eosinophils was more pronounced in the AD samples. In addition, through the association analysis of immune cells with the identified 17 AD_Inflammatory_MDD shared genes, we observed that 16 MDD-associated genes had different degrees of positive or negative correlations with infiltrating immune cells in the AD samples (please refer to Fig.6b for the specific relationships and associated immune cell types).

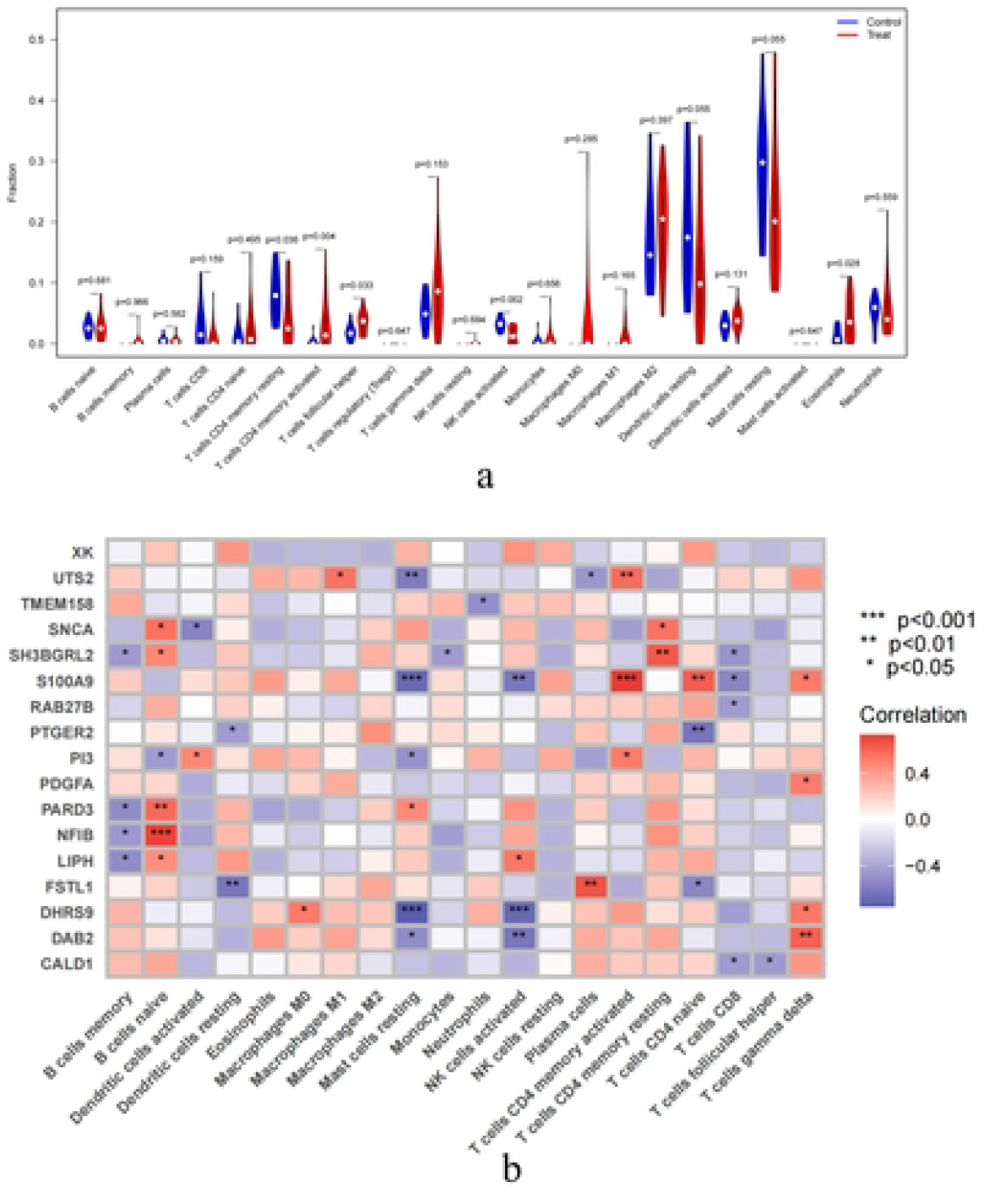

### 3.5 Prediction of key genes by different machine learning models

We assessed the importance of candidate biomarkers by analyzing the results (Fig.7a) of three different machine learning models (Generalized Linear Model - GLM, Random Forest - RF and Support Vector Machine - SVM). The GLM linear model identified PARD3, PDGFA, S100A9, SNCA, and CALD1 as key genes that play a key role in immune response and candidate markers involved in inflammatory response and inflammatory infiltration. RF modeling analysis showed the high importance of SNCA, PARD3, RAB27B, CALD1, and PDGFA in AD and MDD. This was consistent with the results of the GLM, which emphasized the importance of SNCA, PARD3 and RAB27B as possible significant markers. These findings further supported the importance of these biomarkers in the immunoinflammatory process of AD and MDD. The SVM model analysis also revealed the importance of PTGER2, SNCA, CALD1, DAB2, and PDGFA in AD and MDD. The SVM model specifically emphasized the importance of PTGER2 and DAB2, which differed somewhat from the results of the other models, highlighting the different weights assigned to the importance of the biomarkers by the different machine learning methods. However, the three models have a better fitting ability to make predictions to some extent about the key biomarkers of the immune-inflammatory response in the comorbidities of AD and MDD (Fig. 7b, 8c,8d).

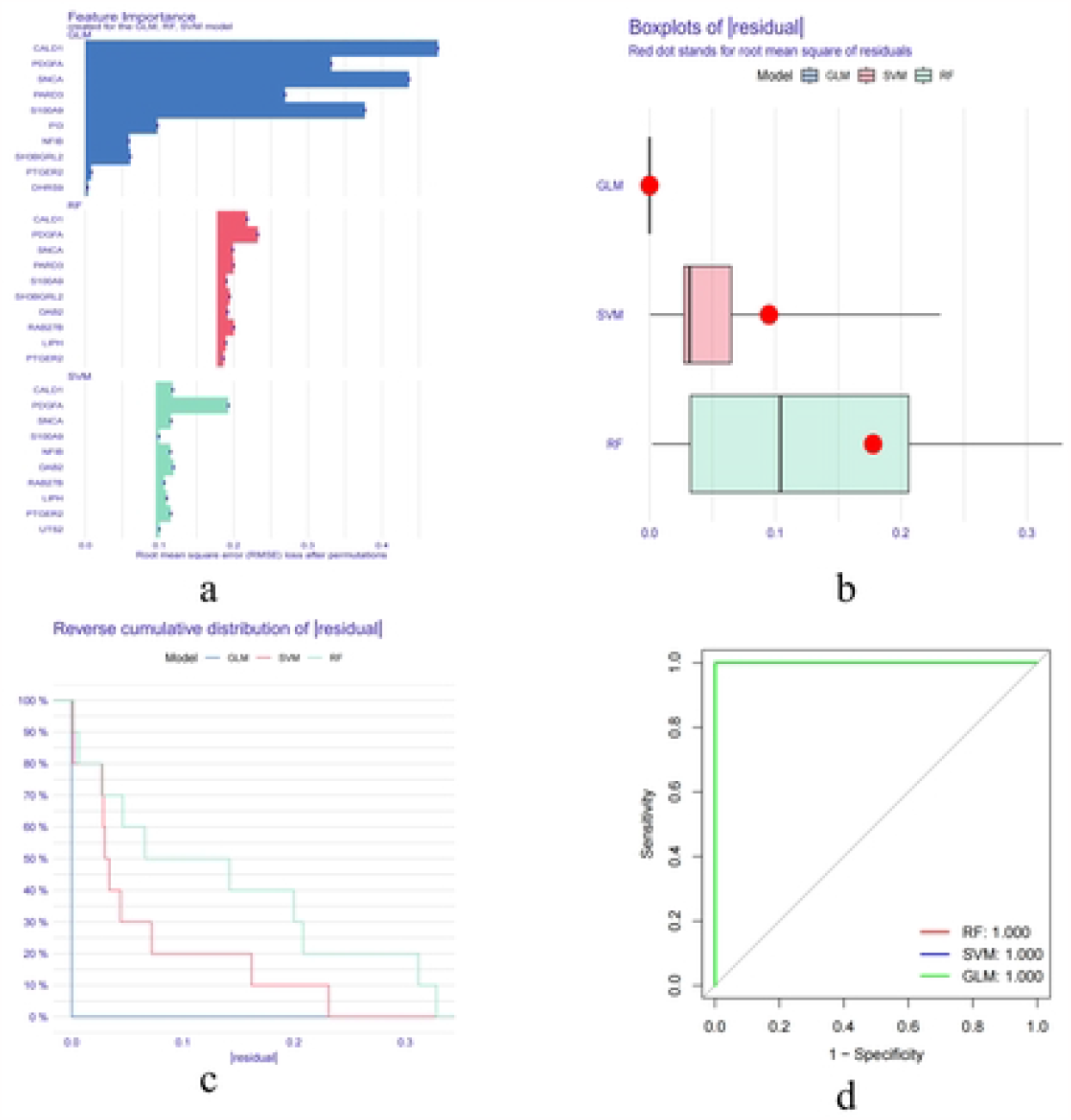

## 4. Discussion

In this study, we integrated the transcriptomes of AD and MDD) and, for the first time, combined comorbidity models with shared genes related to inflammation. This allowed us to unveil common mechanisms between the comorbidities of AD and MDD and the inflammatory responses, further elucidating potential hub genes, shared pathways, and relevant immune cells. Among the shared AD_Inflammatory_MDD genes, we identified 9 hub genes (SNCA, S100A9, SH3BGRL2, RAB27B, TMEM158, DAB2, FSTL1, CALD1, and XK). These genes are likely involved in biological processes such as positive regulation of secretion by cell, positive regulation of secretion, positive regulation of proteolysis, and synaptic vesicle recycling, which are possibly mediated by DAB2, RAB27B, SNCA, and S100A9. Immunoinfiltration analysis of the shared genes revealed the presence of 22 types of immune cells, with 16 genes exhibiting correlations with infiltrating immune cells. In the machine learning algorithms, PARD3, PDGFA, S100A9, SNCA, and CALD1 were identified by the GLM model as candidate biomarkers playing key roles in immune inflammatory responses and inflammation infiltration. The RF model identified SNCA, PARD3, RAB27B, CALD1, and PDGFA, while the SVM model pinpointed PTGER2, SNCA, CALD1, DAB2, and PDGFA. These findings provide potential research targets for studying AD and MDD comorbidities.

We conducted a functional enrichment analysis of the 17 shared genes involved in Biological Processes (BP), Cellular Components (CC), Molecular Functions (MF), and KEGG pathways. Notably, we observed significant enrichment in biological processes related to the regulation of mesenchymal cell proliferation, mesenchymal cell proliferation, and glial cell differentiation. The skin is one of the primary tissues affected by AD. Research has demonstrated that AD patients commonly experience chronic inflammation and aberrant immune responses in their skin, resulting in symptoms such as itching, redness, and dry skin. The regulation of mesenchymal cell proliferation appears to be pertinent to skin pathophysiology in AD because mesenchymal cells are integral components of the skin and play a role in cell proliferation and tissue repair(Na et al., 2014). Moreover, there could be irregularities in the proliferation or function of Mesenchymal Stem Cells (MSCs) in AD, potentially compromising the skin barrier. A compromised skin barrier can heighten the sensitivity of the skin to external stimuli, further exacerbating the inflammatory responses(Kim et al., 2023). While there are a few studies on depression focusing on the regulation of mesenchymal cell proliferation, an emerging body of research indicates that mood disorders may not only impact the brain but could also be linked to systemic inflammatory responses(Qiao, Xu, Zhao, Ye, & Zhang, 2008; Yamanashi et al., 2017). The increased levels of immune cells and inflammatory factors in individuals suffering from depression can potentially influence physiological processes throughout the body, including cell proliferation. Specifically, irregularities in mesenchymal cell proliferation may cause neuroinflammation, a pivotal characteristic of depression(Skok et al., 2022). AD frequently presents with compromised skin barrier function, rendering the skin more susceptible to external stimuli. Mesenchymal cells are instrumental in the repair and regeneration of the skin, including the restoration of the damaged stratum corneum(Song et al., 2023). Hence, mesenchymal cell proliferation may be associated with repairing and rejuvenating the skin barrier in individuals with AD. Furthermore, an intricate relationship exists between MDD and the inflammatory responses of the immune system(Farid Hosseini et al., 2007). Elevated inflammatory factors are associated with the development and severity of MDD. Mesenchymal cells may influence the development and symptom severity of depression by suppressing the inflammatory responses or modulating immune cell activity(Siddiqui et al., 2023; Xiong, Mahmood, & Chopp, 2024). Thus, mesenchymal cell proliferation may be associated with the development of MDD by influencing the immune-inflammatory responses. Glial cell differentiation involves the differentiation of cells, such as astrocytes, precursor cells of hematopoietic stem cells, and oligodendrocytes that support neurons(Gonzalez-Barriga et al., 2021). This biological process plays a key role in the central nervous system (CNS) and is primarily involved in neuronal protection, support, and regulation. Previous studies elucidated a complex interplay between the immune and nervous systems, commonly referred to as immune-neural regulation(Maximova et al., 2021). During this regulation, astrocytes and microglia can generate and release inflammation-related cytokines, potentially affecting neuronal activity and function(Loveland, Yu, Churilov, Yassi, & Watson, 2023). This immune-neural regulation may exhibit abnormalities in individuals with AD and MDD, impairing the nervous system function(X. Huang, Li, & Wang, 2023).

The KEGG analysis of shared genes revealed significant enrichment in pathways such as Neuroactive ligand-receptor interaction, IL-17 signaling pathway, and Rap1 signaling pathway. Neuroactive ligand-receptor interaction is a biological signaling pathway intricately linked to the functioning of the nervous system. It encompasses interactions between the surface receptors of nerve cells and neurotransmitters, hormones, or other neuroactive substances(Y. Yang et al., 2023). This signaling pathway is critical in numerous physiological processes, including nerve conduction, neural development, and the regulation of emotions. Imbalances in neurotransmitters and neuroactive receptors are widely shown to be linked to depression. Consequently, neuroactive ligand-receptor interaction significantly affects mood regulation and depression pathogenesis(L. Chen, Fu, Du, Jiang, & Cheng, 2023). The IL-17 signaling pathway is closely tied to the immune system and is implicated in various autoimmune and inflammatory diseases(Navarro-Compan et al., 2023). IL-17 is an inflammatory cytokine initially linked to autoimmune diseases like psoriasis and rheumatoid arthritis(Saviano et al., 2023). Nonetheless, recent studies unveiled that IL-17 also has a crucial role in AD, impacting the skin keratinization process and resulting in abnormal differentiation of epidermal cells(Balakirski, Burmann, Hofmann, & Kreuter, 2023). Anomalous activation of the IL-17 signaling pathway may cause interactions between immune cells and neurons, subsequently impacting mood regulation and cognitive function. This aligns with the neurobiological underpinnings of depression(Meyer-Arndt et al., 2023). The Rap1 signaling pathway is a significant cellular signaling pathway critical in processes such as cell adhesion, cell proliferation, and cell differentiation(Shelton et al., 2021). In neurons, the Rap1 signaling pathway is involved in synapse formation and plasticity, critical for nervous system function(Kermath et al., 2021). It has been shown that synaptic plasticity and aberrant signaling between neurons are crucial mechanisms in MDD development(Deng et al., 2023). Therefore, irregularities in the Rap1 signaling pathway may be linked to neurobiological changes in MDD.

The enrichment analysis of hub genes suggested the potential significance of DAB2, RAB27B, SNCA, and S100A9 in the pathogenesis of AD and MDD. DAB2 (Disabled homolog 2) is an intracellular signaling protein renowned for its involvement in several biological processes and diseases. Previous studies showed that DAB2 plays a key role in intracellular signaling, cell adhesion and endocytosis(Yu et al., 2015). Currently, there is a dearth of research on the association between DAB2 and AD and MDD. This underscores the significance of exploring the potential relationship between DAB2 and skin cell adhesion, inflammation, and neuronal signaling. Nonetheless, RAB27B (Ras-related protein Rab-27B) is a member of the Rab family of proteins, typically associated with intracellular membrane transport and secretion. Research indicates that RAB27B might regulate immune cell secretion, particularly the release of cytokines and intermediates(Cheng et al., 2020). Aberrant secretion of immune cells can potentially initiate skin inflammation, implying the potential involvement of RAB27B in this process. SNCA (α-synuclein) proteins are intricately tied to the nervous system and are recognized for their association with neurological disorders like Parkinson’s disease. While SNCA is typically linked to nervous system diseases, increasing evidence suggests its potential role in the immune system. Specifically, SNCA expression may regulated during immune inflammation(Xu et al., 2022), and changes in SNCA expression have been associated with severe depression disorder. Nevertheless, the precise pathogenic mechanism of SNCA in depression remains elusive. S100A9 (S100 calcium-binding protein A9) is a calcium-binding protein that has been shown to play a key role in the activation of immune cells and the release of inflammatory mediators(Almeida et al., 2023). A previous study indicated that S100A9 and the upstream gene Nax jointly regulate the inflammatory response of skin tissue. This is associated with the impairment of the epidermal barrier in specific dermatitis, suggesting that S100A9 may be a potential target for treating specific dermatitis(Zhao et al., 2020). S100A9 is also considered a novel and effective regulator of brain inflammation because it can mediate inflammation in microglia/macrophages, ultimately leading to MDD(Sun et al., 2022). These results suggest that as an inducer of inflammation, S100A9 can participate in the pathogenic mechanisms of both AD and MDD.

The increased inflammation infiltration is one of the characteristics of dermatitis, and it is also associated with comorbid MDD in many inflammatory diseases, such as coronary artery disease and co-occurring diabetes and obesity(Maekawa, Sugimoto, Kume, & Ohta, 2022). We found that increased infiltration of the resting memory CD4 T cells, activated memory CD4 T cells, follicular helper T cells, activated NK cells, and eosinophils was more pronounced in AD samples. These findings demonstrate the complexity of the immune system in the AD pathogenesis. The increase in these cell types may be related to the inflammatory and immune responses in the skin of AD patients. Moreover, there was a marked increase in the infiltration of different subtypes of CD4+ T cells in AD, which may be related to skin inflammation and immune response(Inaba et al., 2023). Early childhood atopic dermatitis is typically associated with high IgE levels. Increasing evidence suggests that follicular helper T cells (Tfh) are crucial in promoting IgE production. In this study, the immunoinfiltration analysis showed a significant increase in Tfh (Jiang et al., 2021). Natural killer cells (NK) are associated with the pathophysiology of AD, and immune cell analysis demonstrated that NK cells and T cell subpopulations are associated with the severity of AD(Worm et al., 2023). The increase in eosinophils may be related to the inflammation and allergic characteristics of AD because they typically participate in allergic reactions and tissue inflammation(Naito & Kumanogoh, 2023). Interestingly, research has also found significant immune infiltration in MDD(S. Huang, Li, Shen, Liang, & Li, 2023). These findings provide a foundation and reference for future exploration of the association between AD and MDD.

There was consistency in the results of the different machine learning models, with SNCA and PARD3 being identified as important biomarkers in the three models. The alpha-synuclein protein encoded by SNCA plays a crucial role in neurotransmission and neuroplasticity. Therefore, changes in SNCA expression could be associated with MDD. Research has suggested that SNCA may contribute to depression pathogenesis through complement-mediated microglial phagocytosis and inflammatory responses, thus leading to synaptic loss and neuronal cell death in the hippocampus(Du et al., 2021). These shared inflammatory responses provide a potential mechanism for investigating the comorbidities of AD and MDD. PARD3 is an essential cell polarity protein, and its high importance value likely reflects its critical role in maintaining cell-cell communication and signal transduction(Valdivia et al., 2020). PARD3 may be involved in cellular polarity and synaptic formation among neurons, which is crucial for effective communication and signal transmission between neurons(Schmidt, Mariconda, Morillo, & Apraku, 2014). Although the specific role of PARD3 in skin diseases remains unclear, it may play an important role in maintaining cell polarity and barrier function. Specific dermatitis is often associated with the disruption of skin barrier function and inflammatory infiltration, and the high importance value of PARD3 suggests that it may contribute to these processes.

### Limitation

This study utilized publicly available data, which may be influenced by sample sources and experimental designs, and therefore, caution should be exercised in interpreting the results. Although the machine learning models identified potential biomarkers, further experimental validation of their exact roles in AD and MDD is required. Additionally, AD and MDD are complex conditions influenced by multiple factors, and while immunoinfiltration analysis was conducted, a deeper exploration of their functions and interactions was not undertaken. Thus, in-depth research could contribute to a better understanding of their roles in these diseases.

## 5. Conclusion

This study successfully integrated transcriptomic data of AD and Major MDD using a comorbidity model and machine learning algorithms to unveil potential shared mechanisms between AD-MDD comorbidities and inflammatory responses. SNCA, S100A9, SH3BGRL2, RAB27B, TMEM158, DAB2, FSTL1, CALD1, and XK emerged as pivotal genes in the occurrence of comorbidities between AD and MDD. Immunoinfiltration analysis revealed a significant increase in the infiltration of different subtypes of CD4+ T cells, suggesting a possible association between skin inflammation and immune responses. Moreover, the three machine learning models consistently identified SNCA and PARD3 as important biomarkers.

